# Sub-optimal Discontinuous Current-Clamp switching rates lead to deceptive mouse neuronal firing

**DOI:** 10.1101/2020.08.13.250134

**Authors:** Marin Manuel

## Abstract

Intracellular recordings using sharp microelectrodes often rely on a technique called Discontinuous Current-Clamp to accurately record the membrane potential while injecting current through the same microelectrode. It is well known that a poor choice of DCC switching rate can lead to under-or over-estimation of the cell potential, however, its effect on the cell firing is rarely discussed. Here, we show that sub-optimal switching rates lead to an overestimation of cell excitability. We performed intracellular recordings of mouse spinal motoneurons and recorded their firing in response to pulses and ramps of current in Bridge and DCC mode at various switching rates. We demonstrate that using an incorrect (too low) DCC frequency leads not only to an underestimation of the input resistance, but also, paradoxically, to an artificial overestimation of the firing of these cells: neurons fire at lower current, and at higher frequencies than at higher DCC rates, or than the same neuron recorded in Bridge mode. These effects are dependent on the membrane time constant of the recorded cell, and special care needs to be taken in large cells with very short time constants. Our work highlights the importance of choosing an appropriate DCC switching rate to obtain not only accurate membrane potential readings but also an accurate representation of the firing of the cell.

**Significance Statement:** Discontinuous Current-Clamp is a technique often used during intracellular recordings in vivo. However, incorrect usage of this technique can lead to incorrect interpretations. Poor choice of the DCC switching rate can lead to under- or over-estimation of the cell potential. In addition, we show here that sub-optimal switching rates lead to an overestimation of the cell excitability.

## 1. Introduction

Neurons, by virtue of their plasma membrane and the numerous ion channels that can be found therein, behave—to a first approximation—like RC circuits. Consequently, a stationary electrical (ionic) current flowing through the membrane causes a change of voltage proportional to the resistance of the cell. This is Ohm’s law: V = I × R, where V is sometimes called voltage drop or IR drop. When performing intracellular recordings with microelectrodes, or whole-cell recordings using patch electrodes, electrophysiologists can control the current flowing through their electrode (“current clamp”) to change the membrane potential of the cell and thereby study its excitability. However, the electrode itself, because of its very small tip, acts as an additional RC circuit, and therefore also experiences an IR drop when current is applied. In these conditions, it is essential to be able to separate the physiological response of the cell from a change of voltage caused by the resistance of the very electrode used to perform the recording. Two main techniques have been developed over the years to overcome this problem. The first one, the so-called “Bridge” mode, consists (broadly speaking) in subtracting the voltage drop caused by the current injection through a variable resistor set to a value close to the estimated electrode resistance from the voltage measured by the electrode. This technique works well if the resistance of the electrode can be assumed to be constant over a large range of current intensity. Unfortunately, that is often not the case, particularly with small intracellular microelectrodes, which can exhibit strong nonlinearities (Purves, 1981). A second technique was invented in the early 1970s, which consists in injecting the current and measuring the potential at separate times, hence the name “discontinuous current clamp” (DCC) (Brennecke and Lindemann, 1971; Finkel and Redman, 1984). Instead of injecting a continuous current, the amplifier will alternate at a high frequency between injecting a pulse of current (scaled appropriately so as to conserve the same charge transfer) for a very short duration (classically 1/3 of the DCC period), while no current is injected for the remainder of the DCC period. The membrane potential is sampled at the end of the period when no current is injected through the microelectrode. If the time constant of the electrode is fast enough compared to the time constant of the membrane, then the IR drop through the electrode has had time to vanish when the potential is sampled, while the IR drop through the membrane would have barely decayed. In theory, these two recording modes (Bridge and DCC) should yield the same values of membrane potential, as long as they are used in the proper conditions. One important aspect parameter is the DCC switching rate, which needs to be high enough so that the membrane time constant can smooth out the short pulses of current, but not so high as to prevent the IR drop through the electrode to vanish before the end of the sampling period. An incorrectly set DCC rate should, in theory, only lead to under-or over-estimating the membrane potential. Yet, a recent study (Jensen et al., 2020) illustrates that the firing behavior of a spinal motoneuron in response to a triangular ramp of current can change drastically depending on the DCC switching rate set by the experimenters, suggesting that the choice of the DCC switching rate is a critical parameter to take into consideration not only to obtain accurate readings of the membrane potential but also when studying the firing rates of the cell. In this paper, We demonstrate that using a sub-optimal (too low) DCC frequency lead not only to an underestimation of the cell resistance, but also, paradoxically, to an artificial overestimation of the firing of these cells: neurons fire at lower current, and at higher frequencies than at higher DCC rates, or than the same neuron recorded in Bridge mode.

## 2. Methods

### 2.1. Animals

All procedures were approved by the Paris Descartes University ethics committee (CEEA34; authorization number 2018052100307589) and followed the European Directives (86/609/CEE and 2010-63-UE) and the French legislation on the protection of animals used for scientific purposes. Three C57BL/6 and four B6SJL male mice (weight 25–31 g; 27.9±2.3 g; N=7) were used in this study.

### 2.2. Experimental procedure

The surgical procedures have been described previously (Manuel et al., 2009; Manuel and Heckman, 2012). Briefly, atropine (0.20 mg/kg; Aguettant) and methylprednisolone (0.05 mg; Solu-Medrol; Pfizer) were given subcutaneously at the onset of the experiment, to prevent salivation and edema, respectively. Fifteen minutes later, anesthesia was induced with an intraperitoneal injection of sodium pentobarbitone (70 mg/kg; Pentobarbital; Sanofi-Aventis). A tracheotomy was performed, and the mouse was artificially ventilated with pure oxygen (SAR-830/AP ventilator; CWE). The end-tidal CO2 level was maintained around 4% (MicroCapstar; CWE). The heart rate was monitored (CT-1000; CWE), and the central temperature was kept at 37°C using an infrared heating lamp and an electric blanket. A catheter was introduced in the external jugular vein, allowing us to supplement the anesthesia whenever necessary (usually every 20–30 min) by intravenous injections (sodium pentobarbitone, 6 mg/kg). The adequacy of anesthesia was assessed on lack of noxious reflexes and the stability of the heart rate (usually 400–500 bpm) and end-tidal PCO2. A slow intravenous infusion (50 μL/h) of a 4% glucose solution containing NaHCO3 (1%) and gelatin (14%; Plasmagel; Roger Bellon) helped maintain the physiological parameters. The animal was paralyzed after the surgery with atracurium besylate (Kalceks; initial bolus was 0.1 mg, followed by a continuous infusion 0.01 mg/h). Additional doses of anesthetics were then provided at the same frequency as before the paralysis, and adequacy of anesthesia was assessed on the stability of the heart rate and PCO2. The vertebral column was immobilized with two pairs of horizontal bars (Cunningham Spinal Adaptor; Stoelting) applied on the Th12 and L2 vertebral bodies, and the L3–L4 spinal segments were exposed by a laminectomy at the Th13–L1 level. The Triceps Surae nerve (containing the branches innervating the Medial Gastrocnemius, the Lateral Gastrocnemius, and the Soleus) was dissected and placed on a bipolar electrode for stimulation. All other branches of the sciatic nerve were cut. The tissues in the hind limb and the spinal cord were covered with pools of mineral oil. At the end of the experiments, animals were killed with a lethal intravenous injection of pentobarbitone (200 mg/kg).

### 2.3. Electrophysiological recordings

The motoneurons were impaled with microelectrodes (tip diameter, 1.0–1.5 μm) filled with either 3 M KCl or 2 M K-Acetate (resistance 23.1±5.9 MΩ [16.0–33.0 MΩ], N=13). Recordings were performed using an Axoclamp 2B amplifier (Molecular Devices) connected to a Power1401 interface and using the Spike2 software (CED). The current (Im) and voltage output (10Vm) of the amplifier were low-pass filtered at 10 kHz and sampled at 20 kHz. When recorded, the continuous output I1 and V1, and the DCC monitor output were sampled at 100 kHz. After impalement, identification of motoneurons rested on the observation of antidromic action potentials in response to the electrical stimulation of their axon in the triceps nerve. All care was taken to compensate for the microelectrode resistance and capacitance. No bias current was used to maintain the resting membrane potential. All cells kept for analysis had a resting membrane potential more hyperpolarized than −50 mV and an overshooting antidromic spike. As fully described previously (Manuel et al., 2009), the input resistance was measured using the peak response of a series of small-amplitude square current pulses (−3 to +3 nA, 500 ms) recorded in DCC mode (8 kHz). The membrane time constant was measured on the relaxation of the membrane potential after injection of short hyperpolarizing current pulses (−5 nA, 1 ms), recorded in Bridge mode. Slow triangular ramps of current were injected in DCC mode (switching rates as described in the text) and in Bridge mode when possible. The order in which the different recording modes and DCC rates were applied was randomized in each cell. A recovery period of at least 30s was left in between each repetition. Using an offline automated script, the timing of each spike was recorded along with the current intensity at that time to construct the F-I curve. At switching rates <3 kHz, the DCC voltage trace was often too distorted to identify spikes reliably. In these cases, the continuous voltage trace was carefully scanned manually to identify spikes. The onset current was defined as the value of the injected current at which the first action potential was generated on the ascending phase of the ramp. The offset current was the current intensity corresponding to the last action potential on the descending phase of the ramp. The F-I gain was measured as the slope of the F-I relationship in the most linear part of the ascending phase of the ramp (“primary range”). The voltage threshold was measured at the point when the slope of the membrane voltage crosses 10 V/s (Sekerli et al., 2004) just before the first spike of the ascending phase of the ramp.

### 2.4. Numerical simulations

Numerical simulations were conducted using the Brian2 simulator (v.2.4.1) in Python v.3.8 and using the SciPy ecosystem (v.1.5.0; Virtanen et al., 2020). For investigating membrane potential ripples (Figures 2 and 4), both the cell and the electrodes are modeled as passive RC circuits with equations:

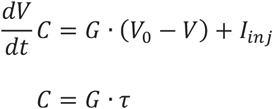

For the cell, Gin was set to 0.2 μS and *τ_m_* to 5 ms. To model the IR drop through the electrode, parameters were chosen so that the electrode was 200× faster than the membrane time constant (τ_e_=*τ_m_*/200=25 μs). The equation above was solved with G_e_=1 μS. Although not quite realistic, this value was chosen so that the response of the electrode would not completely dominate the graphs in Figure 2. Note however that the value of the resistance of the electrode is only relevant at high DCC rates when the IR drop through the electrode does not have time to vanish by sampling time (Figure 2). At lower switching rates, the resistance of the electrode is irrelevant since its contribution has fully dropped to zero at the end of the DCC period. For investigation of the effect of the DCC rate on firing, we used as simple integrate-and-fire model with a passive leak conductance and an after-hyperpolarization (AHP) current (Meunier and Borejsza, 2005; Manuel et al., 2006). The membrane potential (*V_m_*) is governed by the equations:

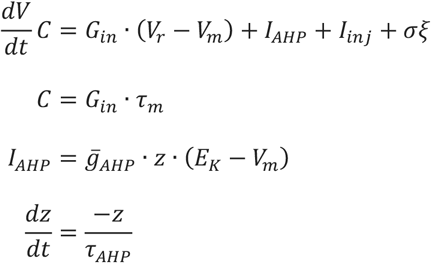

*G_in_* is the input conductance of the cell (we used the values 0.2 μS, 0.4 μS, and 0.67 μS for S, FR, and FF motoneurons, respectively, see text). *τ_m_* is the membrane time constant (varied between 2 and 5 ms, see text). *V_r_* is the resting membrane potential (0 mV). *σ* is a noise term. *I_AHP_* is the AHP current. 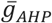 is the maximum conductance of the AHP (2 μS), *E_k_* is the reversal potential of the AHP (−5 mV). *z* is the fraction of the AHP conductance open at any point in time, and *τ_AHP_* is the relaxation time constant of the AHP (10 ms). For simplicity, the dynamic of the AHP during the spike is not modeled, and instead, the parameter *z* is incremented instantaneously at each spike (elicited when *V* > *V_th_, V_th_* =10 mV) according to *z_after_* = (1 – *α*) · *z_before_* + *α*, where *α* is the fraction of the AHP recruited by a single spike (α=0.25) (Meunier and Borejsza, 2005), *z_before_* is the value of *z* just prior to the spike and *z_after_* the value of *z* just after the spike. *I_inj_* is the current injected by the amplifier in Bridge mode. In DCC mode, this current is chopped and scaled with a duty cycle of 1/3. DCC rates range from 1 to 8 kHz.

### 2.5. Code availability

All figures were drawn using matplotlib v.3.2.2 (Hunter, 2007). The code for analysis and production of figures is available at https://doi.org/10.5281/zenodo.4139701

## 3. Results

### 3.1. Case study

Let us start by observing the effect of changing the DCC rate on the response of a motoneuron to a triangular ramp of current. Figure 1 shows a typical example.

**Figure 1.**
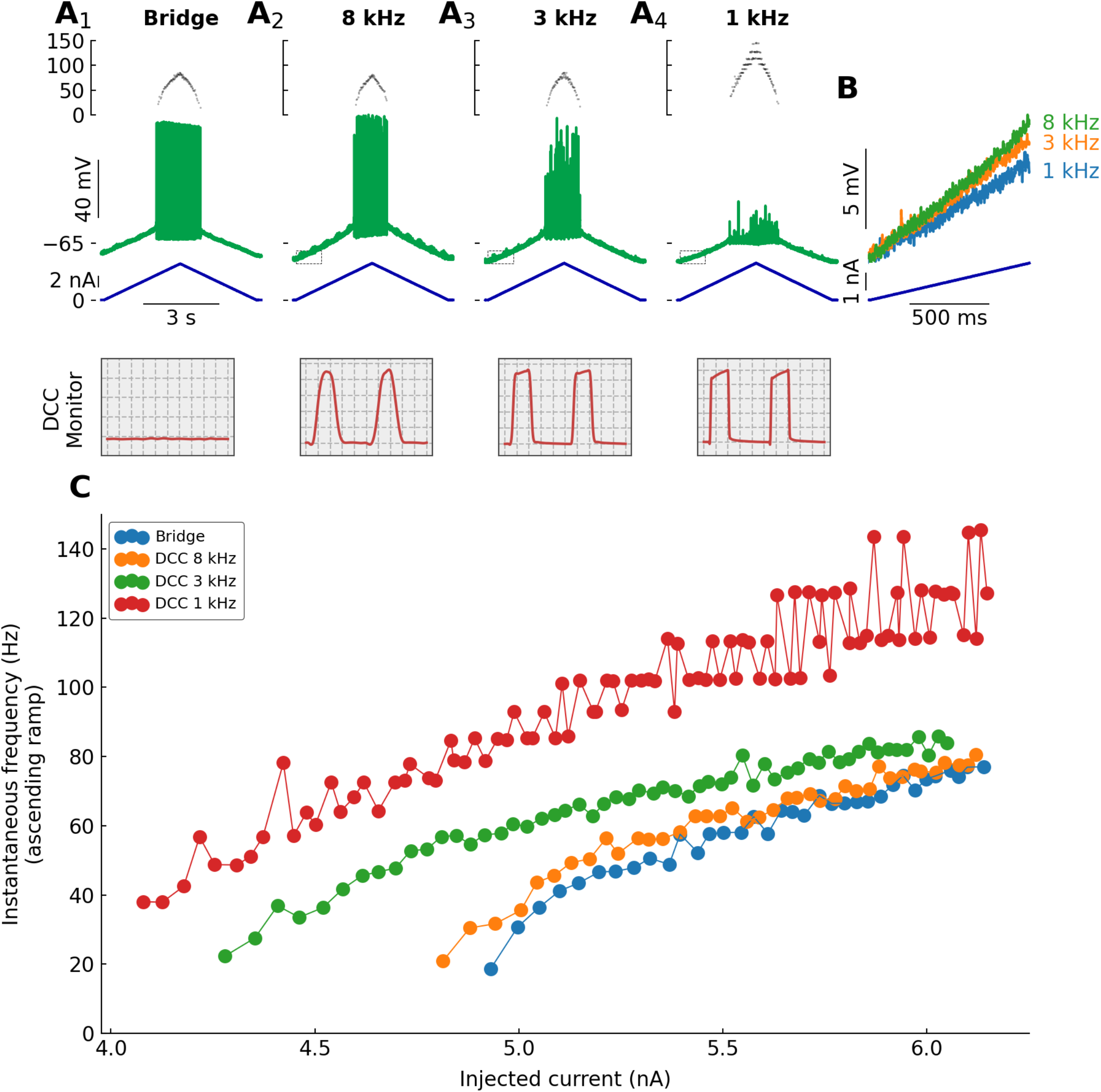
Typical example showing how DCC rates alter the response of a motoneuron to a slow ramp of current. **A.** Response of a triceps Surae motoneuron (*R_in_*=3.9 MΩ; *τ_m_* =4.2 ms) to a slow (2 nA/s) triangular ramp of current, recorded in Bridge mode and in DCC mode with switching rates 8, 3 and 1 kHz. Bottom traces: injected current. Middle traces: voltage response. Top traces: instantaneous firing frequency. The boxes on the bottom represent the monitoring traces used to check the settling of the electrode, recorded at the top of the ramp. Time bases from left to right: 25 μs, 25 μs, 66 μs, and 195 μs per division. **B.** Expansions of the regions delimited with the dashed box in A. **C.** F-I curves showing the instantaneous firing frequency plotted against the injected current at the time of each spike.

Because this motoneuron had a fairly low rheobase and did not require a lot of current to fire, we were able to record the response in Bridge mode. That response is free of artefacts due to DCC switching, and we will therefore use it as the control firing for this cell. In addition, the same ramp of current was injected in DCC mode with switching frequencies 8, 3, and 1 kHz. Before the cell has even started firing, one can already observe a difference in the rate of rise of the voltage at the onset of the ramp (Figure 1B). At 8 kHz, the cell depolarized by 8.2 mV over the first second, while it only depolarized 7.0 mV at 3 kHz and 5.2 mV at 1 kHz over the same period. Since the ramp of current is the same in all cases, this indicates that changing the DCC switching rate affects the apparent resistance of the cell. Note that the trace in Bridge mode is not shown here. With sharp electrodes, it is usually difficult to estimate the resistance of the cell in Bridge mode, as the IR drop through the electrode might change depending on the intensity of the injected current and therefore cannot be perfectly compensated by the Bridge balance circuit. Nevertheless, in all cases, the ramp depolarized the motoneuron progressively until it started to fire repetitively (Figure 1A). The shape of the spikes was strongly affected by lower DCC rates due to the lower sampling rate, and careful manual observation of the trace was required to identify spike times in these conditions. When recorded in Bridge mode or with a DCC rate of 8 kHz, the initial firing was irregular and accelerated very steeply over the first few spikes. Then, the firing became more regular and increased approximately linearly with the injected current (Figure 1C). This is the classical response of mouse spinal motoneurons to this kind of current injected: a brief sub-primary range, followed by a linear primary range (Manuel et al., 2009; Iglesias et al., 2011). The response recorded in DCC mode at 8 kHz is almost indistinguishable from the one recorded in Bridge mode (Figure 1B). However, when recorded with a DCC rate of 3 kHz, although the response was similar, quantitative differences were visible. Paradoxically, even though the apparent resistance of the cell was lower (Figure 1B), the cell started firing at a lower current intensity, and at higher frequencies than at 8 kHz (Figure 1C). These effects were even more pronounced at lower DCC rates. At 1 kHz, the firing started at even lower current intensity, the firing frequency increased very steeply with the injected current and reached much higher values (>100 Hz) than with higher DCC rates. Moreover, when the firing frequency increased beyond ~80 Hz, the firing acquired a very distinctive step-like pattern, where the firing frequency tended to oscillate back and forth between two discrete values (Figure 1C).

### 3.2. DCC switching rate affects the apparent cell resistance

The first effect outlined above, namely the decrease in apparent cell resistance at low DCC rates, is fairly straightforward to explain. By design, in DCC mode, the amplifier injects a short pulse of current, then stops the injection to allow the voltage drop through the electrode to vanish before the membrane potential is sampled. However, during that time of no current injection, the membrane potential will also decay. The technique only works if the electrode time constant (adjusted to be as fast as possible using the capacitance compensation circuit of the amplifier) is much faster than the membrane time constant. In these conditions, the DCC frequency can be set high enough that the membrane potential has barely decayed by the time the voltage is sampled, and the membrane potential recorded in DCC mode is very close to the membrane potential that would be recorded with a perfectly balanced Bridge (Figure 2B). If the DCC rate is too low, however, then the membrane potential has time to decay in between the end of the current pulse and the sampling time (Figure 2A). Consequently, the change in membrane potential recorded in DCC mode is smaller than the true change, thereby producing an underestimation of the input resistance at low DCC rates. Conversely, if the DCC rate is too high, then the IR drop through the electrode does not have time to vanish by the time the potential is sampled (Figure 2C). Therefore the value of the membrane potential of the cell is contaminated by a fraction of the IR drop through the electrode. The change in potential for a given current intensity is larger than expected, thus yielding an overestimation of the input resistance of the cell (Figure 2C). Figure 2D shows how the apparent input resistance changes with the DCC switching rate. At low rates, the contribution of the electrode is completely gone by the time the membrane potential is sampled, and the degree of underestimation of the input resistance is solely dependant on ratio between the DCC switching period and the membrane time constant. At high switching rates, the contamination by the IR drop through the electrode leads to an apparent increase in input resistance. This effect is proportional to the resistance of the electrode (higher resistance electrodes lead to larger overestimations), and it starts at lower DCC rates when the electrode time constant is slower (Figure 2D).

**Figure 2.**
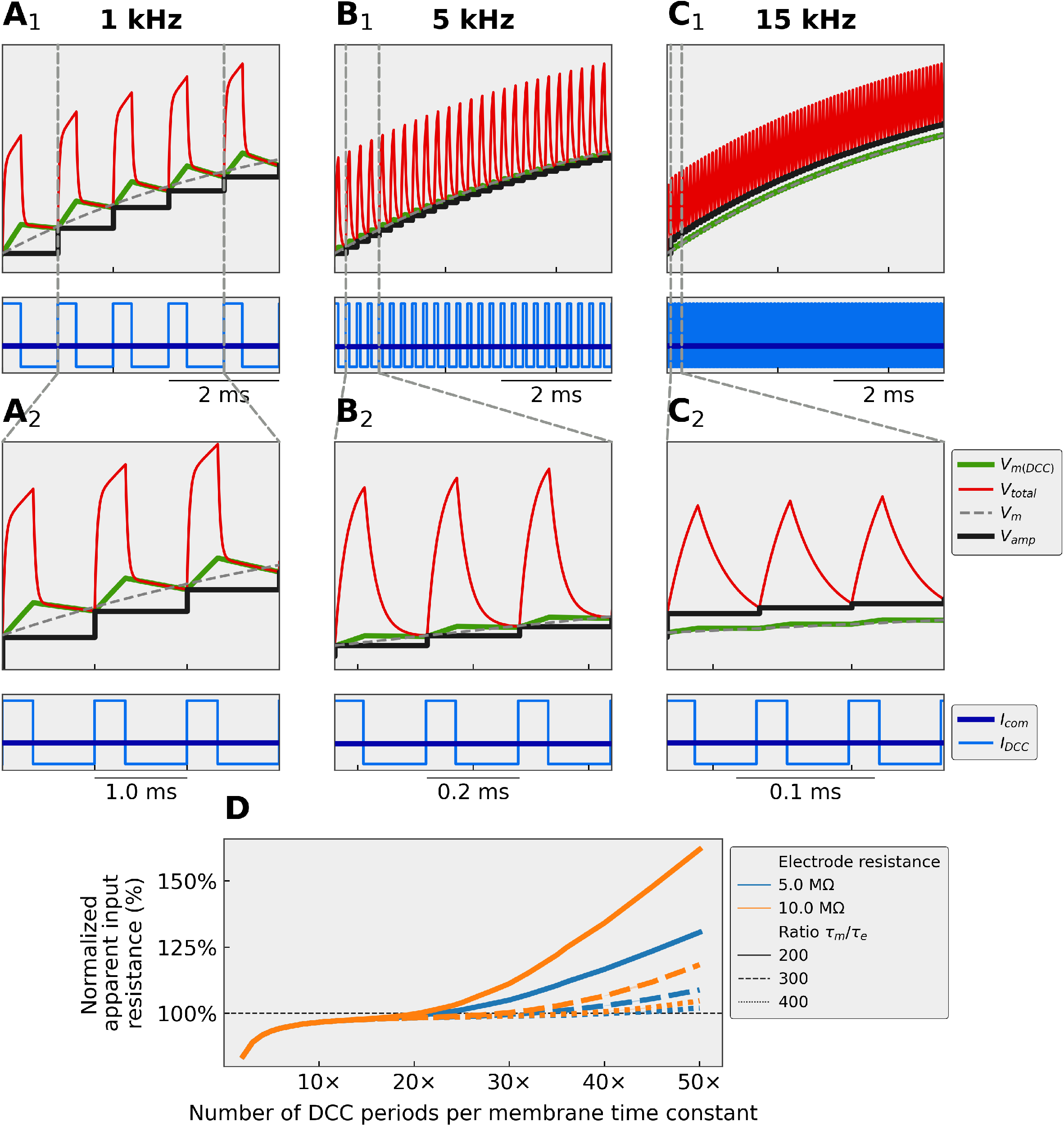
Numerical simulations showing how the DCC rate affects the membrane potential. **A.** recording with a DCC rate of 1 kHz (5 cycles per time constant). **B.** recording with a DCC rate of 5 kHz (25 cycles per time constant). **C.** recording with a DCC rate of 15 kHz (75 cycles per time constant). in A-C, the simulated cell had a resistance of 5 MΩ and a time constant of 5 ms. The electrode had a resistance of 1 MΩ and an effective time constant 200× faster than the membrane time constant. The cell was injected with a 1 nA square pulse of current. *V_m_:* response of the cells to the continuous current as would be observed in an ideal situation where the electrode resistance was perfectly compensated for by the Bridge circuit. *V_total_*: continuous voltage recorded at the tip of the electrode that includes the voltage drop through the electrode and the cell membrane. *V_amp_:* voltage measured by the amplifier in DCC mode, which is the value of *V_total_* sampled at the end of each DCC period and stored in a sample-and-hold circuit. *V_m(DCC)_*: actual potential across the membrane during a DCC injection. This value is not accessible to the experimenter. *I_com_*: stationary current that the experimenter is imposing to the cell. *I_DCC_*: actual current injected in the cell. That current is 3× the amplitude of *I_com_,* but injected for only 1/3 of the time. **D.** Apparent input resistance of the cell (normalized to the real input resistance *R_in_* =2.5 MΩ in this case) as a function of the normalized DCC rate (number of DCC periods per membrane time constant). The measurements were obtained with two different electrode resistances (5 and 10 MΩ) and three different electrode time constants (200×, 300×, and 400× faster than the membrane time constant).

The underestimation of the input resistance at low DCC rates is very consistent across all recorded cells. Figure 3 shows that, in the 13 recorded cells, the apparent input resistance decreases sharply under ~4 kHz. Above that value, the estimated input resistance is relatively constant as the DCC rate increases. The apparent input resistance would start increasing when the IR drop through the electrode does not have time to vanish before sampling time. However, this situation is easily identified on the monitoring scope, and we have therefore not explored higher DCC rates.

**Figure 3.**
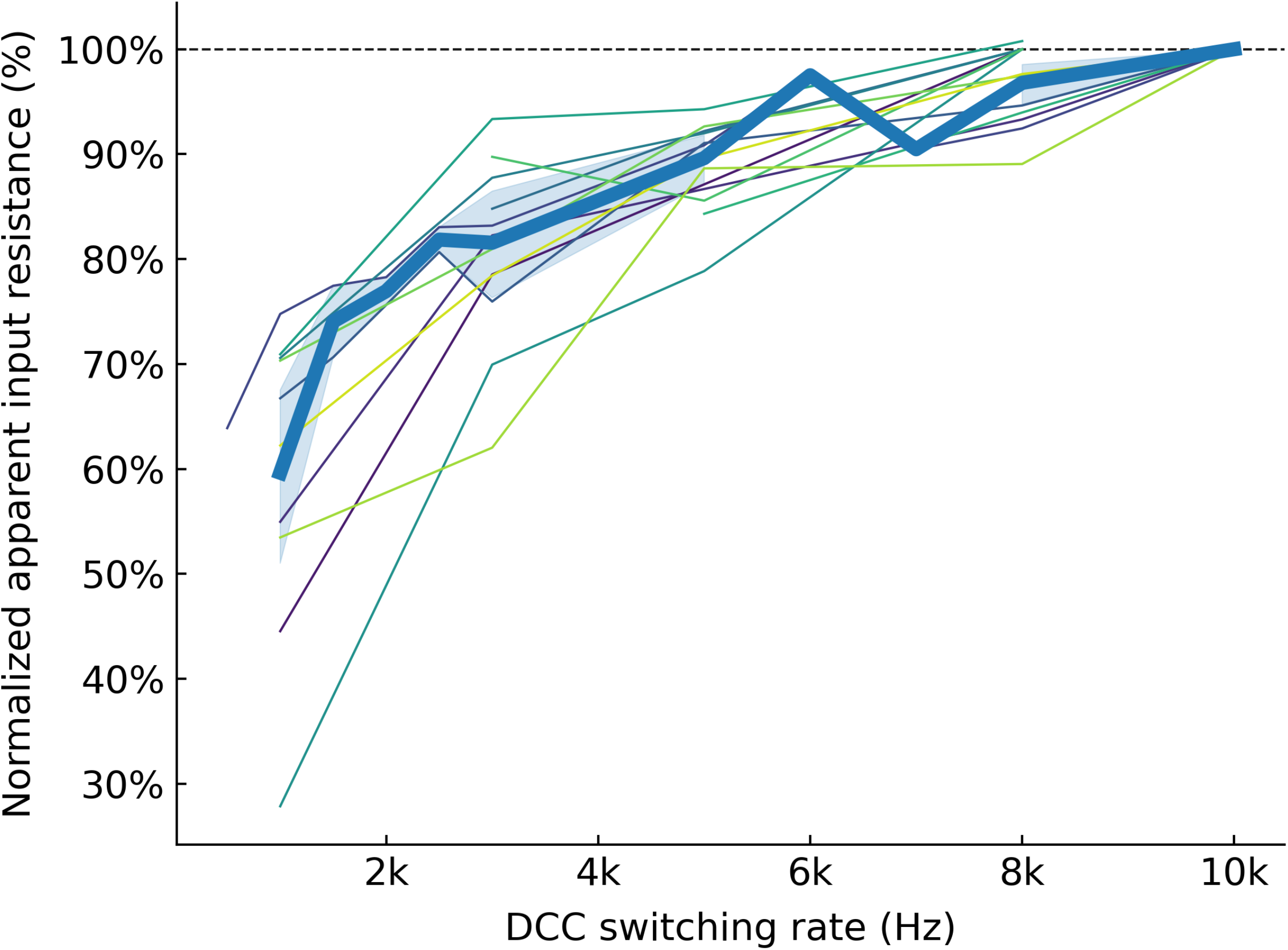
Effect of the DCC rate on the apparent input resistance of the cells. Experimental results showing the apparent cell input resistance as a function of the DCC switching rate. The input resistance of each cell was normalized to the value measured with the highest DCC rate. Each thin line corresponds to a different cell, and the thick line shows the average apparent resistance (±95%CI, shaded area).

### 3.3. Low DCC switching rates can drive firing

The effect of the DCC on the F-I curves is more subtle. Although the amount of charge transferred to the cell is the same in DCC and in Bridge, the frequency content of the input is not the same. By chopping the current injection in short pulses, the DCC introduces harmonics of the DCC frequency in the input signal (Brette and Destexhe, 2012). Moreover, at low DCC rates, the membrane potential has time to increase during the pulse injection and then has time to decay substantially in between each current injection, creating “ripples” in the membrane potential (Finkel and Redman, 1984) (Figure 4). Although these ripples are present in the membrane potential, they are hidden to the experimenter by the DCC sample-and-hold circuit, which samples the potential at the end of the DCC period and holds the amplifier output constant at that value until the next sampling time. We, therefore, relied on numerical simulations to investigate these ripples. Figures 4A_1-3_ show examples of steady-state ripples experienced by a model of a typical FR motoneuron when injected with 10 nA of current in DCC mode at 1, 5, and 15 kHz. Because the actual current injected during the DCC pulses is 3× the intensity of the desired current, these ripples can be quite large. The amplitude of these ripples depends not only on the DCC frequency but also on the time constant of the membrane (as well as, of course, the resistance of the cell and the intensity of the injected current). Figures 4B_1-2_ show how the amplitudes of the ripples change with DCC rate (normalized by the membrane time constant, i.e. number of DCC periods per membrane time constant) for three values of the motoneuron input resistance (1.5 MΩ, 2.5 MΩ, and 5 MΩ, corresponding to typical values for, respectively, FF, FR, and S mouse motoneurons (Martínez-Silva et al., 2018)), and two values of injected current (5 and 10 nA, which are values that are typically reached when injecting ramps of current in mouse motoneurons). These figures show that the amplitudes of the ripples increase steeply when the DCC switching rate is decreased, particularly under 10 DCC periods per time constant. However, even for reasonable rates (10-20 DCC periods per time constant), the ripples can reach several millivolts in amplitude. The net effect of the DCC is therefore a series of (potentially large amplitude) membrane potential ripples (which are hidden by the sample-and-hold circuit of the amplifier), superimposed to the slow depolarization of the quasi-stationary ramp. The spiking observed in these conditions is caused by this mixed dynamic and stationary input, rather than a response to the stationary input alone. The ripples are a very potent stimulus for triggering firing, much more than the slow static depolarization, and as a consequence, the cell fires at lower current and higher frequency when the DCC rate is low compared to when the DCC rate is high or when the cell is recorded in Bridge mode (Figure 1C). This effect is seen consistently across motoneurons.

**Figure 4.**
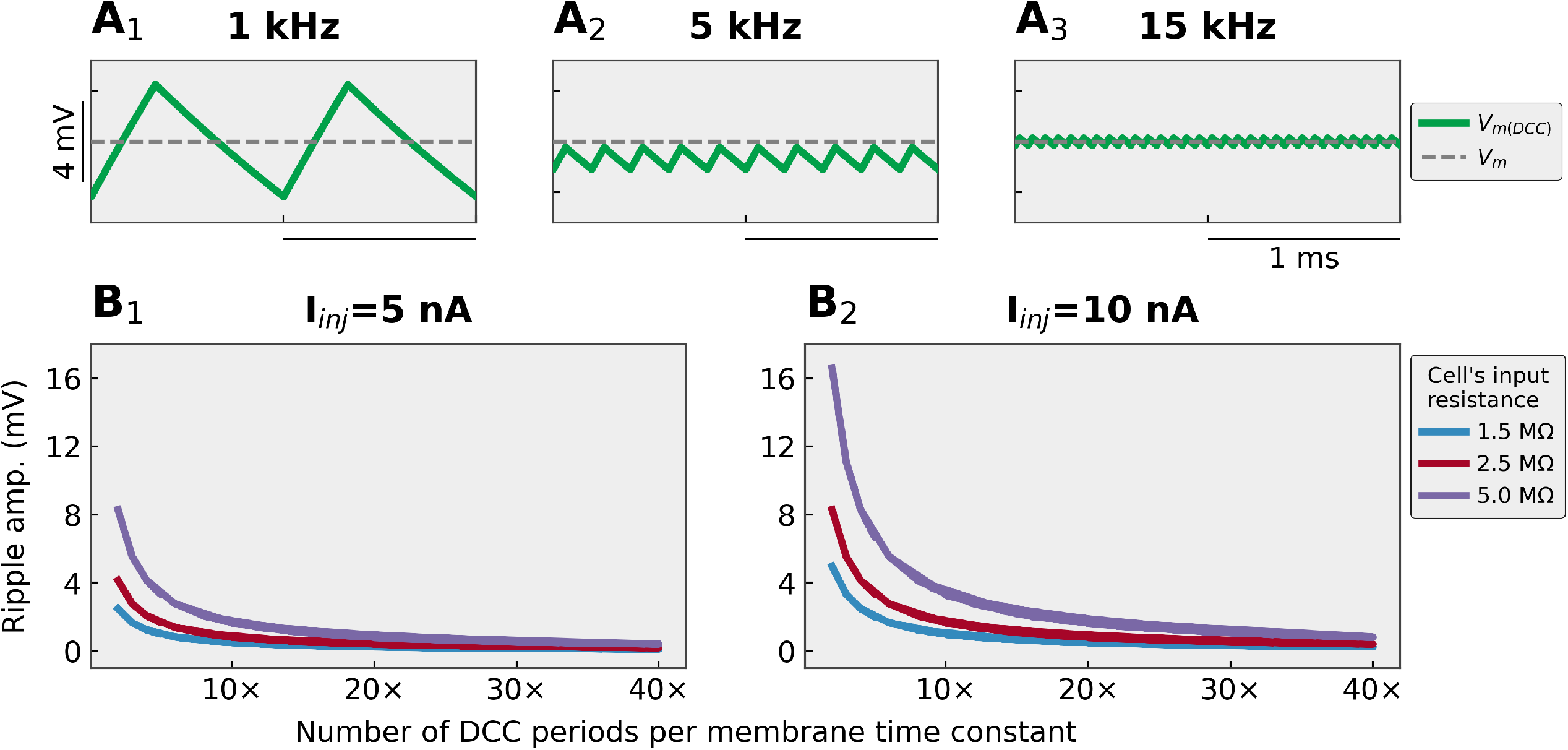
DCC recording mode induces ripples in the membrane potential. **A.** Traces showing the steady-state amplitude of the membrane potential ripples in a cell with an input resistance of 2.5 MΩ, time constant 3 ms, and injected current 10 nA when recorded in DCC mode at 1 kHz (A_1_), 5 kHz (A_2_), and 15 kHz (A_3_). **B.** Plots showing the amplitude of the ripples as a function of the normalized DCC frequency (number of DCC cycles per membrane time constant). The response was measured in steady-state for two current intensities that are routinely reached during our recordings in mouse spinal motoneurons (5 nA B_1_, and 10 nA B_2_) and for three values of the cell’s input resistance 1.5 MΩ, 2.5 MΩ, and 5 MΩ (which correspond to typical values for FF, FR, and S motoneurons, respectively).

Figure 5A–D shows how the current intensity required to start firing (onset current), the current intensity when the cell stopped firing (offset current), the slope of the ascending phase of the F-I relationship (F-I gain), and the voltage threshold measured on the first spike of the ramp vary with DCC frequency. It is apparent that values measured at low DCC rates are usually very different than the ones measured at higher rates, and that the values tend to converge to a stable value when the DCC rate is increased past a critical point. Moreover, in the motoneurons in which we were able to record the response in Bridge mode (Figure 1), the values measured with the highest DCC rates are indistinguishable from the values recorded in Bridge mode (paired mean differences: onset current 0.0963 nA [95%CI −0.111, 0.531], N=9; offset current 0.116 nA [95%CI −0.124, 0.584],N=9; F-I gain −1.04 Hz/nA [95%CI −2.24, 0.0418],N=9), except for the voltage threshold (paired mean difference 8.59 mV [95%CI 3.21, 15.8],N=9), which is expected since the voltage threshold cannot be measured accurately in Bridge mode because of the IR drop through the electrode that may not be fully compensated in this mode. Moreover, These curves demonstrate that the rate of 10 cycles per time constant recommended in the Axoclamp manual is not high enough. Rates of at least 15–20 cycles per time constant are necessary to get good estimates of the value of most of these measurements.

**Figure 5.**
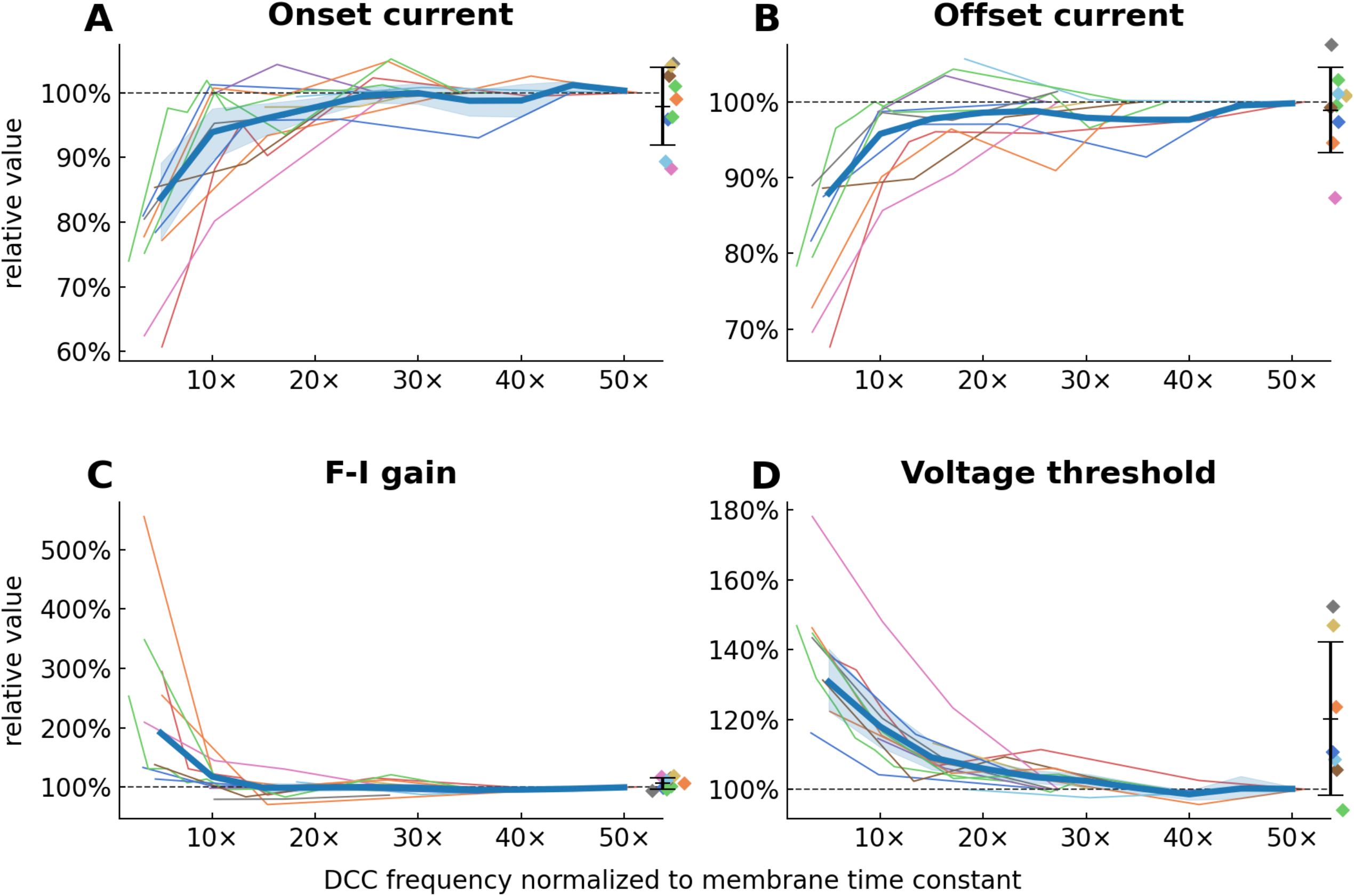
Relationship between parameters measured on the F-I curves and the DCC frequency used during the recording. In all panels, each line represents one motoneuron. Onset current: current required to elicit the first spike on the ascending ramp. Offset current: current at which the firing stops on the descending ramp. F-I gain: slope of the F-I relationship measured on the ascending part of the ramp. Voltage threshold: voltage measured at the foot of the first spike elicited on the ascending ramp. **A-D.** Value of each of the parameters normalized to the value measured at the highest DCC rate achieved in each motoneuron (dashed horizontal line) plotted against the DCC rate normalized by the time constant of each motoneuron. The thick line represents the average values across motoneurons (±95%CI, shaded area). The diamonds on the right side of each plot represent the measurement obtained in Bridge mode (mean ± SD).

### 3.4. Low DCC rates entrain firing at discrete intervals

As shown above, using a low DCC switching rate not only leads to cells firing at lower current but also at higher frequencies. For instance, in the cells exemplified in Figures 1 and 6A, lowering the DCC rate from 8 to 3 kHz led to both a leftward and upward shift of the F-I curve. Reducing it further to 1 kHz led to the appearance of marked “plateaus” in the instantaneous firing frequency. This behavior can be reproduced in a simple integrate-and-fire model (Figure 6B). These plateaus correspond to interspike intervals (ISI) that are multiples of the DCC switching period. The cell no longer fires at its natural interspike interval, but instead is driven to fire on the crest of the membrane potential ripples when the after-hyperpolarization from the preceding spike has relaxed sufficiently for the membrane potential to come close to the voltage threshold (Figure 6C). Because a significant amount of current has to be injected in spinal motoneurons to reach the firing threshold, the ripples can get quite large (Figure 4B), which is why they can entrain firing with shorter ISI (higher frequency) than what would be observed for the same current intensity in Bridge mode (Figure 6C).

**Figure 6.**
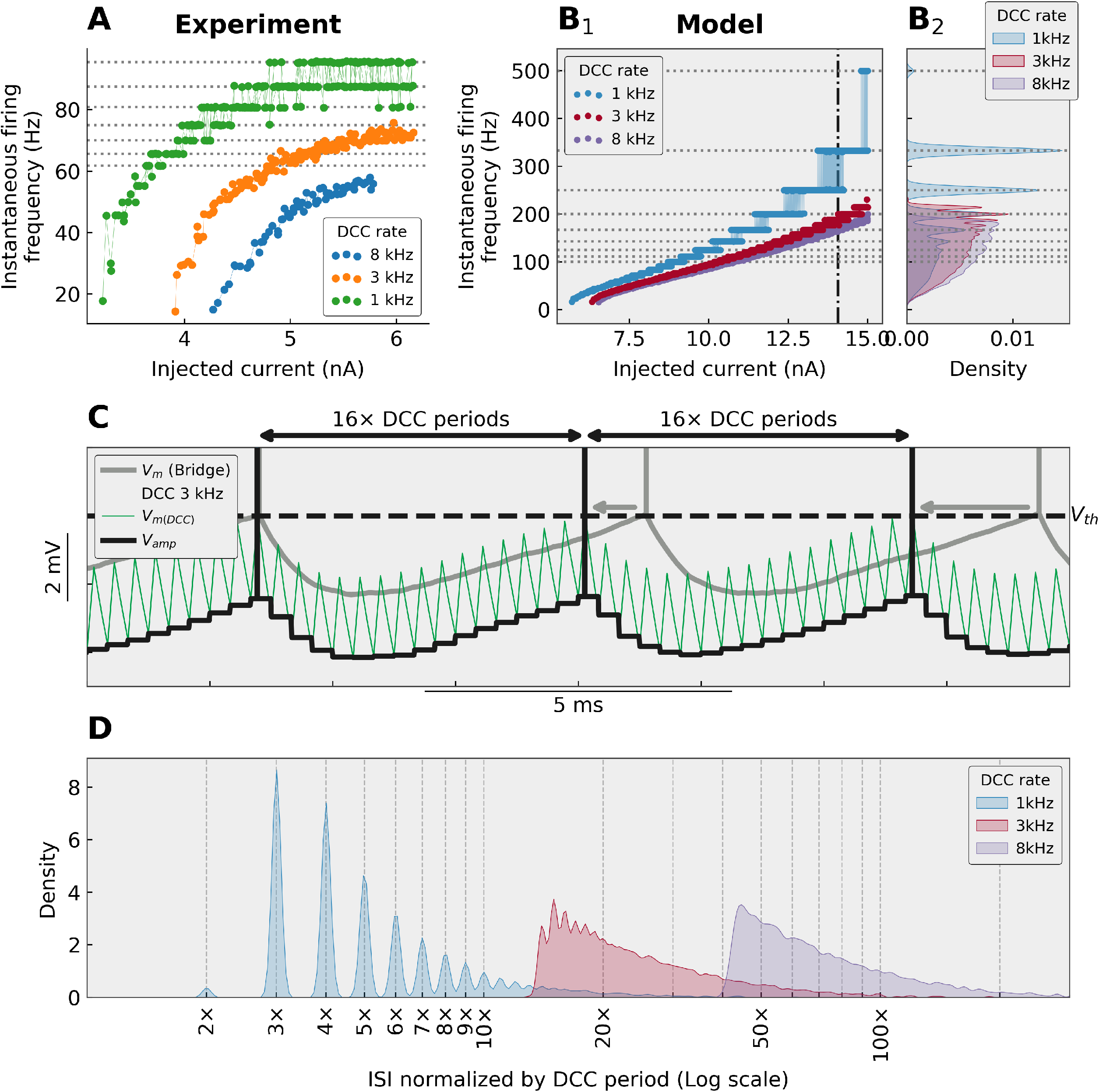
Stepwise pattern is a sign of sub-optimal DCC rate. **A.** F-I curves from a Triceps Surae motoneuron (*R_in_*=2.0 MΩ; *τ_m_*=3.8 ms), injected with a triangular ramp of current (1 nA/s) and recorded in DCC mode at three different DCC switching rates. At low DCC rates, a clear stepwise pattern is apparent, which corresponds to multiples of the switching rate (1050 Hz in this instance): 95.5 Hz (or 1 spike every 11 DCC periods), 87.5 Hz (1:12), 80.8 Hz (1:13), 75.0 Hz (1:14), 70.0 Hz (1:15), 65.6 Hz (1:16), etc. **B.** The same phenomenon can be observed in a simple integrate-and-fire motoneuron model. The model was that of a typical FF motoneuron (*R_in_*=1.5 MΩ; *τ_m_*=2.0 ms), injected with a 10 nA slow ramp of current (1 nA/s), and recorded in DCC mode at 8, 3, and 1 kHz. A stepwise pattern is apparent at the top of the F-I curve at 3 kHz and is evident at 1 kHz (see distinct peaks in the distributions of the firing frequencies in B_2_). The horizontal dotted lines represent the multiples of the period of the 1 kHz switching rates. The vertical dash-dotted line represents the region zoomed-in in C. **C.** Comparison of the behavior of the model recorded in Bridge (grey line) and DCC mode at 3 kHz. The thick black line represents the Vamp output of the amplifier, while the thin green line represents the true membrane potential *V_m(DCC)_* which is hidden from the experimenter by the sample-and-hold circuit. The membrane potential ripples created by the DCC shorten the interspike intervals (grey arrows) and entrain the firing with interspike intervals that are multiples of the DCC period. **D.** Distribution of the interspike intervals obtained in DCC mode at 1, 3, and 8 kHz. The intervals have been normalized by the DCC period (1 ms, 0.33 ms, and 0.125 ms, respectively) and plotted on a logarithmic scale. At 1 kHz, the interspike intervals are concentrated at multiples of the DCC period.

The plateaus are characteristic of recordings with sub-optimal DCC rates for two reasons. Firstly, the amplitude of the ripples decreases with increasing DCC rates (Figure 4), therefore they are less likely to “stick out” from the noise and entrain firing. Secondly, the plateaus are only apparent when the firing rate of the cell approaches the fundamental frequency of the DCC. Consider the behavior of the model in Figure 6B with a sub-optimal DCC frequency of 1 kHz (2 DCC cycles per membrane time constant). Firing starts at a low frequency then increases linearly without visible plateaus until the frequency reaches 50-60 Hz where they are barely visible but become much more prominent above 100 Hz. In this case, the plateaus appear when the firing is entrained at about one spike every 10 DCC cycles, and become more and more prominent as the firing frequency gets closer to the DCC rate: the distribution of the interspike intervals becomes more and more peaked at multiples of the DCC period (Figure 6D). Below 20 DCC cycles, even if entrainment happens, the difference between being entrained at one spike per e.g. 30 or 31 DCC cycles is drowned in the variability of the discharge. Therefore, at higher DCC frequencies, not only do the ripples become smaller, but the range of firing frequencies over which plateaus are apparent is pushed higher and higher, often beyond the normal range of firing frequencies of neurons attainable by neurons. For instance, in the model (Figure 6B), even though the firing frequencies largely overlap when comparing DCC rates of 1, 3, and 8 kHz, firing rates reach one spike every 15 DCC cycles at 3 kHz (plateaus are clearly visible, Figures 6B_1-2_, 6D), but barely reach one spike every 45 DCC cycles at 8 kHz (Figure 6D), and no plateaus are visible (Figure 6B). Plateaus can be observed in all recorded motoneurons (N=13), but not necessarily at the same DCC rates. Plateaus are sometimes visible when recording in DCC mode with a switching rate of 3 kHz if the firing frequency is high enough (Figure 7A). However, even if no plateaus are visible at 3 kHz because the firing rate is too low, further reducing the DCC switch rate systematically leads to the eventual appearance of plateaus as the DCC rate gets closer to the firing rates explored by the cell. Plateaus were never observed with DCC rates of 8 kHz or higher (N=13).

**Figure 7.**
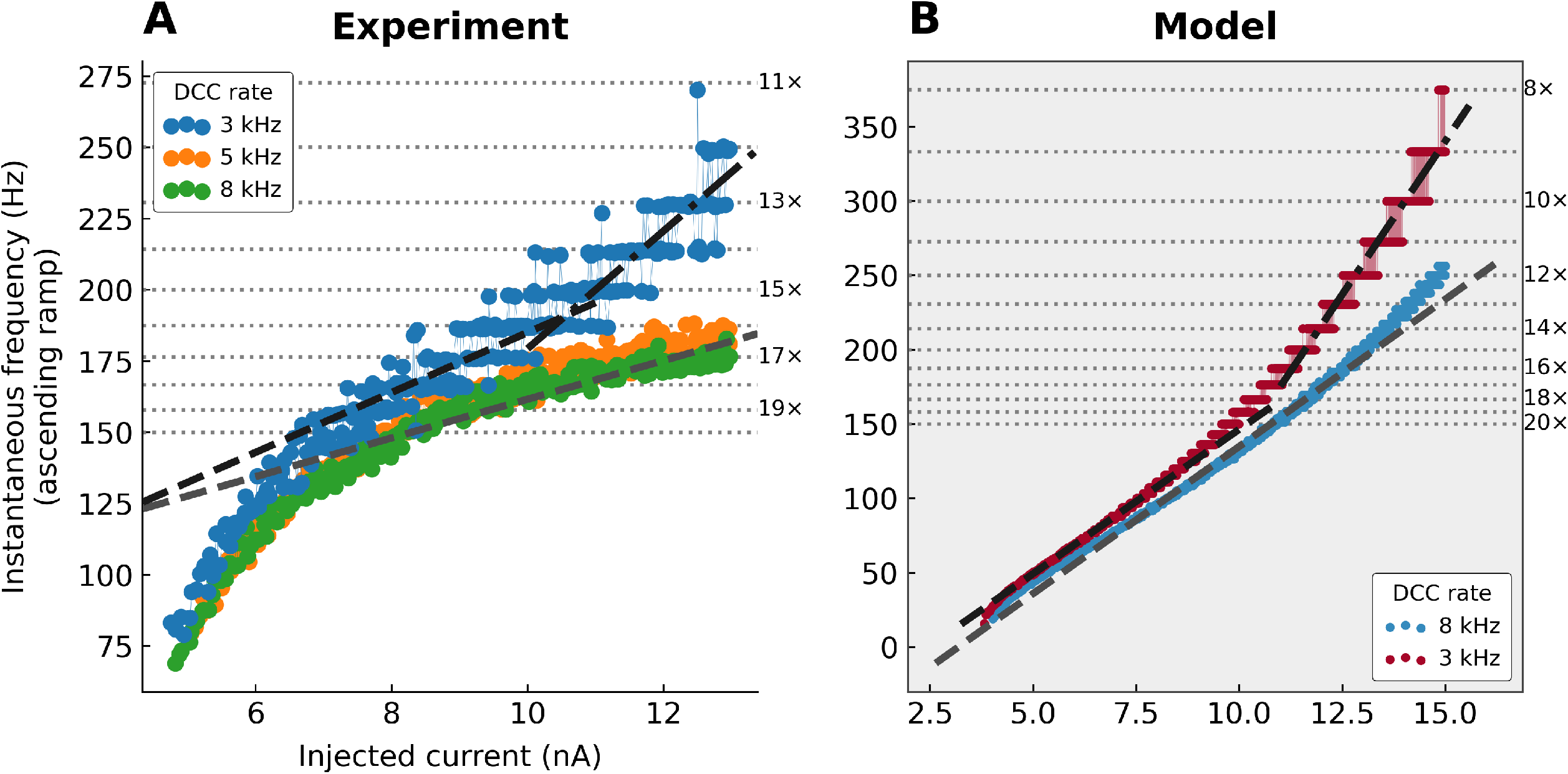
Apparent change of slope in the F-I curve associated with discrete firing intervals. **A.** F-I curves from a Triceps Surae motoneuron (*R_in_*=4.1 MΩ; *τ_m_*=3.3 ms). A fast triangular ramp of current (amplitude 13 nA, 5 nA/s) was injected to drive the firing at high frequency. The instantaneous firing frequency is plotted against the ascending ramp current intensity. Grey dashed line: slope of the F-I curve recorded at 8 kHz measured in the second half of the curve. Black dashed line: slope of the F-I curve recorded at 3 kHz, measured over the range 7–10 nA. Black dash-dotted line: slope of the F-I curve recorded at 3 kHz, measured over the range 11–13 nA. Horizontal dotted lines: subharmonics of the 3 kHz DCC rate. The numbers on the right of each line show the number of DCC period per ISI. **B.** F-I curves obtained in a model with *R_in_*=2.5 MΩ and *τ_m_*=2.0 ms. Compared to the F-I curve obtained with a high DCC rate of 8 kHz, which is mostly linear (grey dashed line), the F-I curve obtained with a DCC rate of 3 kHz changes slope at ~10 nA, from a slope roughly equal to the one measured at 8 kHz to a much steeper slope.

Interestingly, this entrainment effect at sub-optimal DCC rates can induce an apparent increase in the slope in the F-I relationship, particularly when the firing frequency reaches high values. This phenomenon is illustrated in Figure 7 where a motoneuron was stimulated with a high-amplitude (13 nA) fast ramp of current (5 nA/s), expressly for the purpose of pushing the motoneuron to high firing frequencies. At DCC rates 5 and 8 kHz, the resulting F-I curves are almost indistinguishable, with a sub-primary range of current where the frequency was increasing steeply, followed by a region where the frequency increased at a smaller rate (~7 Hz/nA, dashed grey line). When recorded with a DCC rate of 3 kHz, the F-I curve was shifted upward to higher firing frequencies. The slope of the initial linear phase was slightly higher (~10 Hz/nA, black dashed line) than with higher DCC rates. When the firing reached frequencies above 150 Hz, a clear step-wise pattern became apparent and the F-I curve became steeper (~20 Hz/nA), creating the illusion of a “secondary range”, even though this change of slope is not present in the data recorded with higher DCC rates for the same current intensities. This change of slope can be explained by the discretization of the ISI described above. When the DCC frequency is sub-optimal, ISI are entrained at multiple of the DCC period (Figure 6D). As the injected current and the frequency increases, the ISI shorten linearly by discrete steps (e.g. 1 spike every 10 DCC periods, then 1 spike every 9 DCC periods, then every 8 periods, etc.). Since the frequency is the inverse of the ISI, the firing frequency is increasing very steeply as it jumps from plateau to plateau (Figure 7B).

### 3.5. Low DCC rates can trigger firing when cells should not fire

As discussed above, using a low DCC frequency becomes equivalent to injecting a series of short pulses of current. This kind of stimulus is highly efficient to trigger motoneuron firing, much more than a continuous current injection (Delestrée et al., 2014; Martínez-Silva et al., 2018). Figure 8A shows the response of a motoneuron to the same 200 ms-long 4 nA pulse of current, recorded in DCC mode with a rate of 1.5 kHz and 8 kHz. With a DCC at 8 kHz, this pulse of current was not able to reach the firing threshold (Figure 8A_1_). At the lower DCC rate, however, although the amount of current injected is the same, the motoneuron responded with a strong repetitive discharge (Figure 8B_1_). Interestingly, the voltage threshold for the first spike was below the steady-stage potential reached with a DCC of 8 kHz (compare dashed lines in Figure 8A_2_ and B_2_), suggesting that the appearance of firing at 1.5 kHz was not due to a larger depolarization (in fact, the depolarization is smaller, see the effect of the DCC rate on the apparent resistance of the cell, above), but rather due to the strong sensibility of the cell to transient currents and ripples in their membrane potential. In the extreme case, a low DCC rate can turn a motoneuron that was not able to fire repetitively in response to a stationary input (Figure 8C_1_) into a motoneuron that elicits a bout of repetitive firing to the same ramp (Figure 8C_2_). Observation of the DCC monitoring trace on the oscilloscope (inserts in Figure 8C) confirms that the inability to fire repetitively was not due to the electrode becoming blocked. The IR drop through the electrode had fully vanished by the end of the DCC period, and the membrane potential was therefore accurately measured. With a DCC at 1 kHz, however, the DCC period was so long that not only the IR drop through the electrode had time to settle to zero, the membrane potential also rose and decayed during each DCC period.

**Figure 8.**
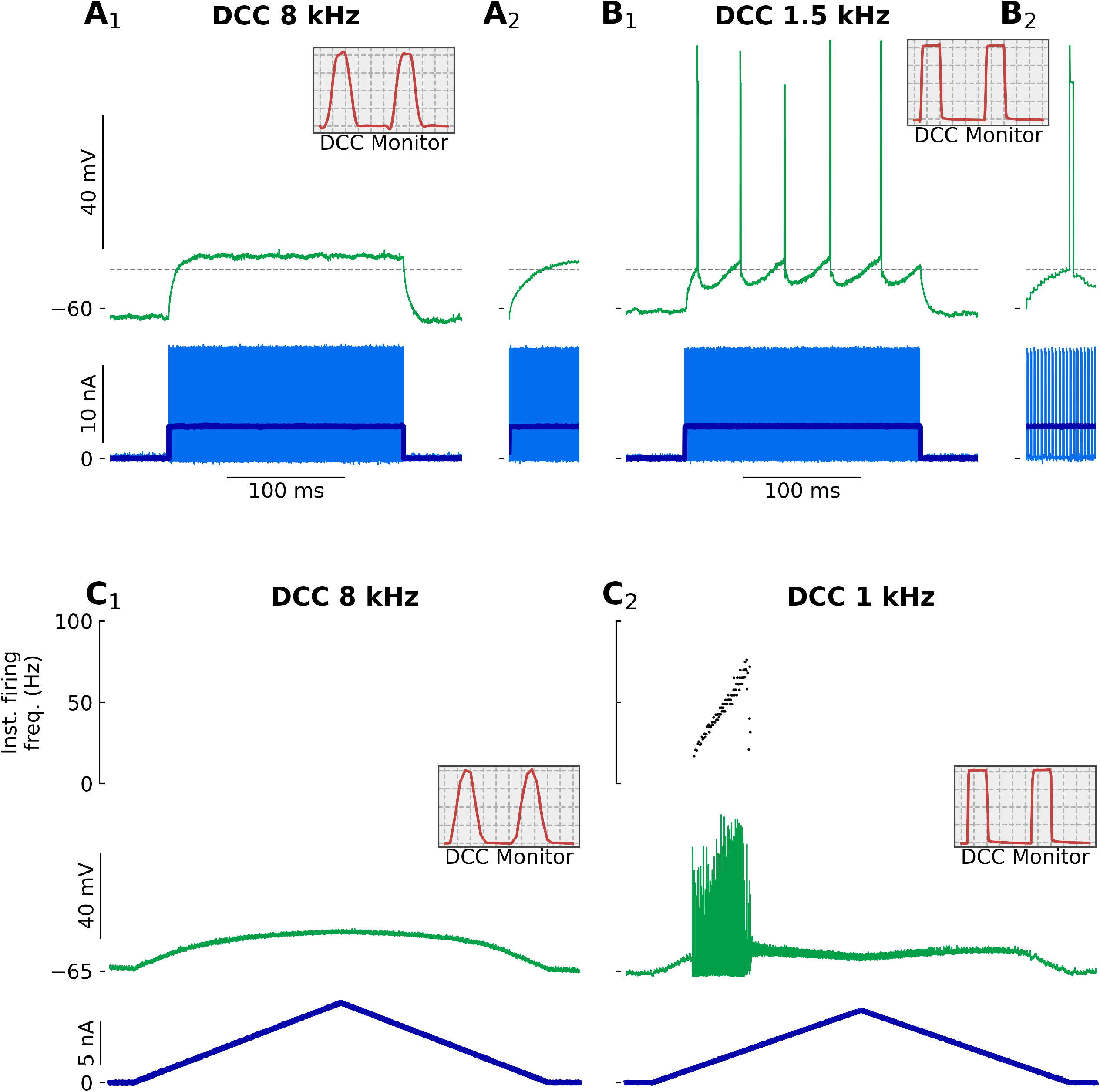
Spurious firing elicited by low DCC rates. **A-B.** Recording from a Triceps Surae motoneuron (*R_in_*=3.5 MΩ; *τ_m_*=4.9 ms) following the injection of a 200 ms-long 4 nA pulse of current. A. Response recorded with a DCC rate of 8 kHz. The inset in A_2_ is a zoom over the first 15 ms following the onset of the pulse. B. Response recorded with a DCC rate of 1.5 kHz. The inset in B_2_ is a zoom over the first 15 ms following the onset of the pulse. The horizontal dashed line represents the voltage threshold measured at the foot of the first spike of the response in B. The grey boxes in A and B represent the monitoring traces used to check the settling of the electrode, recorded at the top of the ramp. Time bases: 25 μs (C_1_), and 133 μs (C_2_) per division. **C.** Response of a Triceps Surae motoneuron (*R_in_*=5.0 MΩ; *τ_m_*=4.7 ms) to the injection of a triangular ramp of current (1 nA/s) with a DCC rate of 8 kHz (C_1_) and 1 kHz (C_2_). The bottom trace is the injected current, the middle trace is the membrane potential and the top graph is the instantaneous firing frequency. The inserts represent the monitoring traces used to check the settling of the electrode, recorded at the top of the ramp. Time bases: 25 μs (C_1_), and 200 μs (C_2_) per division.

## 4. Discussion

This paper describes the effect of an incorrectly set DCC rate on the firing properties of spinal motoneurons. Although a low DCC rate leads to an underestimation of the membrane potential, and therefore an apparent decrease in input resistance, we show that, paradoxically, it has the potential to artificially drive the cells to fire at lower currents and higher frequency, as well as to profoundly alter the shape of the F-I relationship.

### 4.1. Why use the DCC mode?

Given the potential issues outlined above, one might ask what is the advantage of using the DCC mode to record the firing of the cell. First and foremost, the use of the DCC mode is warranted when one wants to obtain a precise measurement of the membrane potential while injecting current through an intracellular microelectrode. Some properties of the neurons (e.g. voltage threshold for firing, trajectory of the membrane potential between spikes) can only be measured in DCC mode. However, if one is only interested in the timing of the spikes, Bridge mode can be sufficient (as demonstrated in some of the measurements in the present study). Unfortunately, sharp microelectrodes often display very strong nonlinearities in the sense that their impedance changes as a function of the intensity of the injected current (Brette and Destexhe, 2012; Purves, 1981). Given the high impedance of the sharp microelectrodes and the amount of current required to elicit firing in spinal motoneurons, the IR drop through the microelectrode often dwarfs the actual membrane potential of the cell. In many cases, at some point on the ascending leg of the ramp, the input stage of the amplifier saturates and it is no longer possible to record the membrane potential at all. The use of the DCC mode can generally alleviate this problem. Because the membrane potential is sampled after the IR drop through the electrode has decayed to a value close to zero, there is less risk that the amplifier would saturate, and spike times can be recorded accurately. Electrode nonlinearities are also associated with an increase in electrode time constant as the intensity of the injected current increases. Thus the chosen DCC rate may be appropriate at the beginning of the ramp when the current is still small, but halfway through the ascending ramp the IR drop through the electrode no longer decays to zero by sampling time, and the voltage cannot be accurately measured. In some extreme cases, the remaining IR drop can be large enough to saturate the amplifier, even in DCC mode. It could be tempting in these conditions to lower the DCC rate to allow more time for the electrode contribution to vanish before sampling time. However, as the present study demonstrates, that could lead to an inaccurate representation of the excitability of the cell. Instead, the safer choice is to discard such recordings and try again with a different electrode, which hopefully will exhibit less nonlinearity.

### 4.2. Practical considerations for the choice of the DCC switching rate

Although there are no theoretical upper limits to the DCC cycling rate, in practice, one is limited by the time constant of the electrode and the capacitance neutralization circuit of the amplifier. That maximum rate can be found by observing the continuous electrode potential on an oscilloscope synchronized to the DCC sampling clock. The goal is to adjust the electrode capacitance compensation circuit and the DCC switching rate to reach the highest DCC rate possible while ensuring that the response shown on the oscilloscope appears flat, that is to say, that the contribution of the electrode resistance to the recorded potential has dropped down to zero before the time when the voltage is sampled. More importantly, there is a lower limit to the DCC rate. Yet it is not always straightforward to know whether the DCC rate is fast enough to not distort the firing of the cell. One reason for this is the fact that the optimal rate depends on the membrane time constant of the cell. A switching rate that is appropriate for one cell might not be optimal for a different cell, or cell type, with a shorter time constant. The first practical consideration for setting the DCC switching rate is therefore to know the time constant of the cells one is recording from. The Axoclamp manual states that the rate must be such “that there are ten or more cycles per membrane time constant. This enables the membrane capacitance to smooth the membrane voltage response to the current pulses” (Axon Instruments, 2003). Our experiments, however, show that this recommendation is too conservative. We show that DCC rates of at least 15–20 cycles per time constant are required to produce measurements that match the ones obtained in Bridge mode (Figure 5). Above this threshold value of 15–20 cycles per time constant, our experiments show that measurements become largely insensitive to the exact DCC rates (Figure 5). One practical application of this result is that one could inject a constant current in DCC mode in a cell while progressively increasing the switching rate starting from a low value (~1 kHz). The depolarization (for positive currents) should start small and increase as the switching rate is increased, then reach a plateau value for a wide range of switching rates, until one reaches the critical rate at which the electrode is no longer fast enough and the depolarization starts to increase again (Figure 2D). The optimal DCC rate is somewhere in the range where the potential plateaus, preferably in the upper end of switching rates, but maybe not exactly the maximum rate to avoid issues linked to the nonlinearities of the electrodes at other current intensities. The fact that, when the switching rate is adequately set, measurements are largely independent of the DCC rate is crucial for electrophysiologists. Indeed, we tend to choose a DCC rate that is close to optimal, and then use the same switching rate for all the cells, despite the fact that they might have slightly different membrane time constants. Thankfully, as long as the switching rate is fast enough, small differences in DCC rates (relative to the membrane time constant) between cells should not impact their respective firing behavior.

### 4.3. DCC mode and neuronal firing

Compared to the Bridge mode, the DCC transforms the input signal from a continuous variation in current intensity to a discontinuous situation, where the current can only be injected as short square pulses. This difference is almost negligible when the DCC rate is high enough for the membrane potential to barely move during the current injection and the subsequent inter-pulse interval. However, at lower DCC frequencies, the membrane potential exhibit substantial ripples (Figure 4). Although these ripples are present across the membrane of the recorded cell, they are hidden from the experimenter by design of the amplifier. The output of the amplifier is held constant at the level of the previous sampled value for the whole duration of a DCC period (thick black line in Figure 2A). When considering slow ramps of current, like in the present study, decreasing the DCC rate from a high frequency to a lower frequency, therefore, amounts to transitioning from a situation where the membrane potential is increasing slowly, to a situation where sharp voltage ripples are superimposed to a slow depolarization. These ripples are particularly efficient at triggering action potentials, particularly when the membrane potential is very close to the firing threshold. It has been shown in many neuronal types that the faster the rate of rise of the membrane potential, the more reliably a spike will be generated (Mainen and Sejnowski, 1995; Azouz and Gray, 2000; Agrawal et al., 2001; Kuo et al., 2006). This high dynamic sensitivity explains why motoneurons recorded with a low DCC rate fire at lower currents despite reaching lower membrane potentials. It also accounts for the fact that, at low DCC rates, firing becomes entrained by the DCC. Membrane potential ripples trigger a spike more reliably than the slow decay of the after-hyperpolarization that follows the preceding spike. The interspike intervals thereby can only take values that are multiples of the DCC period, leading to the characteristic step-like pattern observed in the F-I curves at sub-optimal DCC rates.

### 4.4. The particular case of spinal motoneurons

Interestingly, motoneurons have a natural regime of firing where a similar step-like pattern can be observed in response to slow current ramps. We have shown previously that, in a narrow range of current intensities, motoneurons exhibit subthreshold oscillations, which alternate with spikes, producing a very irregular firing. This regime, called mixed-mode oscillations (MMOs) is responsible for the sub-primary firing range. These oscillations naturally emerge from a Na/K ratio too weak to generate full-blown spikes with high reliability; but when a spike is finally generated, it is phase-locked with oscillations (Iglesias et al., 2011). However, since the frequency of the MMOs is much lower (100–125 Hz, Manuel et al., 2009; Iglesias et al., 2011), and disappear when the firing reaches past the transition frequency between the sub-primary and the primary range (Iglesias et al., 2011), the resulting plateaus in the F-I curve are only apparent over the sub-primary range. In spinal motoneurons, there is a strong relationship between membrane time constant and cell size, such that small, Slow-type (S) motoneurons have a longer membrane time constant than the larger, fast fatigable (FF) motoneurons (Gustafsson and Pinter, 1984). Consequently, FF motoneurons require an even higher DCC frequency than S motoneurons to obtain accurate measurements of their excitability. We have previously shown that mouse motoneurons have shorter time constants than cats (Manuel et al., 2009). Mouse FF motoneurons have an average time constant of 2.1±0.2 ms, FR motoneurons 2.9±0.9 ms, while S motoneurons have a time constant of 4.0±0.7 ms (unpublished data from Martínez-Silva et al., 2018). Based on our present results, which show that a DCC frequency corresponding to at least 15 cycles per time constant is required to measure the excitability of the cell, FF motoneurons should be recorded with a DCC rate of at least 7 kHz, while S motoneurons can accommodate DCC frequencies as low as 3.75 kHz. Because of their size, FF motoneurons are also the cells that require the most current to fire. The impedance of the electrode is often highly non-linear, and both the resistance and the time constant of the electrode tend to increase with the amount of injected current. Consequently, It is often difficult to record the firing of these cells at high DCC rates. Instead, it would be tempting, particularly in these cells, to lower the DCC rate to obtain proper settling of the electrode’s IR drop, but, as we demonstrate here, doing so would lead to an overestimation of the cell’s firing and excitability parameters. Moreover, in a mouse model of Amyotrophic Lateral Sclerosis, we have shown that the largest motoneurons become incapable of firing repetitively in response to a slow ramp of current (Martínez-Silva et al., 2018). Given the membrane time constants of these cells, it was essential to perform these recordings at high DCC rates (all of our recordings were performed in DCC at 7–9 kHz), since lower DCC rates have the potential to distort the firing of these cells, and even mistakenly transform a non-repetitively-firing motoneuron into a repetitively-firing motoneuron (Jensen et al., 2020).

## 5. Conclusions

In conclusion, the effect of inappropriate DCC switching rates on the apparent resistance of the cells is well known. However, the effects on the firing characteristics of the neurons are not often discussed. We show here that choosing a sub-optimal DCC rate may dramatically distort parameters that are classically used to define the “excitability” of neurons: lower current onset, lower current offset, higher firing frequencies, higher F-I gains, and even the appearance of an artifactual “secondary range” of firing. Low DCC rates can therefore lead to a misrepresentation of neuronal excitability.

## Acknowledgements

The author wishes to thank Drs. Daniel Zytnicki, Rob Brownstone, and Marco Beato for their valuable comments and discussions. This work has benefited from the support and expertise of the animal facility and microscopy platform of BioMedTech Facilities at Université de Paris (INSERM US36/UMS2009).

## References

Agrawal N, Hamam BN, Magistretti J, Alonso A & Ragsdale DS (2001). Persistent sodium channel activity mediates subthreshold membrane potential oscillations and low-threshold spikes in rat entorhinal cortex layer V neurons. Neuroscience 102, 53–64.

Axon Instruments I (2003). Axoclamp-2B, Microelectrode Clamp, Theory and Operation.

Azouz R & Gray CM (2000). Dynamic spike threshold reveals a mechanism for synaptic coincidence detection in cortical neurons in vivo. Proceedings of the National Academy of Sciences of the United States of America 97, 8110–8115.

Brennecke R & Lindemann B (1971). A chopped-current clamp for current injection and recording of membrane polarization with single electrodes of changing resistance. T-I-T-J Life Sci. 1, 53–58.

Brette R & Destexhe A (2012). Intracellular recording In Handbook of Neural Activity Measurement. Cambridge University Press.

Delestrée N, Manuel M, Iglesias C, Elbasiouny SM, Heckman CJ & Zytnicki D (2014). Adult spinal motoneurones are not hyperexcitable in a mouse model of inherited amyotrophic lateral sclerosis. The Journal of physiology 592, 1687–703.

Finkel AS & Redman S (1984). Theory and operation of a single microelectrode voltage clamp. Journal of Neuroscience Methods 11, 101–127.

Gustafsson B & Pinter MJ (1984). Relations among passive electrical properties of lumbar alpha-motoneurones of the cat. The Journal of physiology 356, 401–31.

Hunter JD (2007). Matplotlib: A 2d graphics environment. Computing in Science & Engineering 9, 90–95.

Iglesias C, Meunier C, Manuel M, Timofeeva Y, Delestrée N & Zytnicki D (2011). Mixed mode oscillations in mouse spinal motoneurons arise from a low excitability state. The Journal of neuroscience: the official journal of the Society for Neuroscience 31, 5829–40.

Jensen DB, Kadlecova M, Allodi I & Meehan CF (2020). Spinal motoneurones are intrinsically more responsive in the adult G93A SOD1 mouse model of Amyotrophic Lateral Sclerosis. The Journal of Physiology n/a _eprint: https://physoc.onlinelibrary.wiley.com/doi/pdf/10.1113/JP280097.

Kuo JJ, Lee RH, Zhang L & Heckman CJ (2006). Essential role of the persistent sodium current in spike initiation during slowly rising inputs in mouse spinal neurones. The Journal of Physiology 574, 819–834.

Mainen Z & Sejnowski T (1995). Reliability of spike timing in neocortical neurons. Science 268, 1503–1506.

Manuel M & Heckman CJ (2012). Simultaneous intracellular recording of a lumbar motoneuron and the force produced by its motor unit in the adult mouse in vivo. Journal of visualized experiments: JoVE 70, e4312.

Manuel M, Iglesias C, Donnet M, Leroy F, Heckman CJ & Zytnicki D (2009). Fast kinetics, high-frequency oscillations, and subprimary firing range in adult mouse spinal motoneurons. The Journal of neuroscience 29, 11246–11256.

Manuel M, Meunier C, Donnet M & Zytnicki D (2006). The afterhyperpolarization conductance exerts the same control over the gain and variability of motoneurone firing in anaesthetized cats. The Journal of physiology 576, 873–886.

Martínez-Silva MdL, Imhoff-Manuel RD, Sharma A, Heckman C, Shneider NA, Roselli F, Zytnicki D & Manuel M (2018). Hypoexcitability precedes denervation in the large fast-contracting motor units in two unrelated mouse models of ALS. eLife 7, e30955.

Meunier C & Borejsza K (2005). How membrane properties shape the discharge of motoneurons: a detailed analytical study. Neural computation 17, 2383–2420.

Purves RD (1981). Microelectrode methods for intracellular recording and ionophoresis Biological techniques series. Academic Press, London ; New York.

Sekerli M, Del Negro CA, Lee RH & Butera RJ (2004). Estimating action potential thresholds from neuronal time-series: new metrics and evaluation of methodologies. IEEE transactions on bio-medical engineering 51, 1665–1672.

Virtanen P, Gommers R, Oliphant TE, Haberland M, Reddy T, Cournapeau D, Burovski E, Peterson P, Weckesser W, Bright J, van der Walt SJ, Brett M, Wilson J, Jarrod Millman K, Mayorov N, Nelson ARJ, Jones E, Kern R, Larson E, Carey C, Polat ‘I, Feng Y, Moore EW, Vand erPlas J, Laxalde D, Perktold J, Cimrman R, Henriksen I, Quintero EA, Harris CR, Archibald AM, Ribeiro AH, Pedregosa F, van Mulbregt P & Contributors S (2020). SciPy 1.0: Fundamental Algorithms for Scientific Computing in Python. Nature Methods 17, 261–272.

